# A spatially-specific pattern of cortical thickness in frontal lobe predisposes to faster future amyloid accumulation in individuals with subjective cognitive decline

**DOI:** 10.64898/2026.07.11.737919

**Authors:** Diego Vidaurre, Andrea Moreno, Oscar Sotolongo-Grau, Pablo Oyarzo, Juan Pablo Tartari, Angela Sanabria, Montserrat Alegret, Lluís Tárraga, Miren Jone Gurruchaga, Xavier Montalbán, Agustín Ruiz, Mercè Boada, Marta Marquié, Christopher Butler, FACEHBI study group, AMYPAD consortium

**Affiliations:** Centre de Recerca Matemàtica, Barcelona, 08193, Spain; Oxford Centre for Human Brain Activity, Wellcome Centre for Integrative Neuroimaging, Department of Psychiatry, University of Oxford, Oxford, OX3 7JX, United Kingdom; Department of Clinical Medicine, Center of Functionally Integrative Neuroscience, Aarhus University, 8000 Aarhus, Denmark; Department of Biomedicine, Aarhus University, 8000 Aarhus, Denmark; Ace Alzheimer Center Barcelona – Universitat Internacional de Catalunya. Barcelona, Spain; Centro de Investigación Biomédica en Red de Enfermedades Neurodegenerativas (CIBERNED), Instituto de Salud Carlos III, Madrid, Spain; Universitat de Vic-Central de Catalunya (UVic-UCC); Glenn Biggs Institute for Alzheimer’s & Neurodegenerative Diseases, University of Texas Health Science Center, San Antonio, TX USA; Department of Microbiology, Immunology and Molecular Genetics, Long School of Medicine. University of Texas Health Science Center, San Antonio, TX, USA; Department of Brain Sciences, Imperial College London, UK; The George Institute for Global Health, School of Public Health, Imperial College London, UK

## Abstract

The risk of developing AD is conferred through numerous genetic and environmental factors. How these risks influence brain structure prior to the onset of AD pathology is not yet known. We hypothesise that there may be specific anatomic endophenotypes that predispose to AD years before the first signs of amyloid accumulation occur. Using clinical, genetic and brain MRI data from Fundació ACE Healthy Brain Initiative (FACEHBI) —a longitudinal cohort of individuals with Subjective Cognitive Decline aged > 50 years—, we identified a spatially-specific, fine-grained pattern of cortical thickness in frontal lobe regions among amyloid-negative individuals at baseline that holds a significant relationship with the rate of amyloid accumulation during the next five years. This pattern may represent a structural fingerprint of increased susceptibility to the initiation of the AD cascade.

## Introduction

While the genetic and environmental predisposition for Alzheimer disease (AD) is well established (Reitz, Pericak-Vance, Foroud, & Mayeux, 2023; Migliore & Coppedè, 2022), much less is known about how these risk factors are reflected in evolving anatomical markers of vulnerability. Neuropathologic hallmarks of AD such as amyloid-β (Aβ) silently accumulate before overt cognitive symptoms are apparent (Jack, et al., 2013), but what triggers abnormal Aβ accumulation in the first place is not fully understood. It is possible that subtle alterations in brain structure create the conditions that facilitate the initial development of AD neuropathology. This is important, as a neuroimaging-based fingerprint of vulnerability could offer an unprecedented opportunity for prevention and improve our understanding of the earliest stages of AD (Ottoy, et al., 2025).

Longitudinal biomarker studies suggest that brain Aβ accumulation follows a gradual and dynamic trajectory, and that cognitive decline is more strongly associated with Aβ accumulation rate than with Aβ cross-sectional burden (Klinger, et al., 2024). Aβ accumulation can be quantified using the Centiloid (CL) scale derived from amyloid PET imaging (Klunk, et al., 2015). CL values equal or lower than 10 suggest absence of amyloid burden (Amadoru, et al., 2020), whereas values larger than 26 are indicative of significant brain amyloid deposition, consistent with AD pathology and clear predictors of progression to dementia (Hanseeuw, et al., 2021). In parallel, brain MRI studies have identified structural correlates of AD and cognitive decline, including hippocampal atrophy, cortical thinning, and large-scale network alterations (Xia, et al., 2023; Fjell, et al., 2014; Dickerson, et al., 2009). Yet many of these MRI findings have been derived from individuals who already exhibit substantial amyloid pathology or clinical impairment, making it difficult to determine whether the observed anatomical changes precede Aβ accumulation or arise because of ongoing disease processes. Comparatively less attention has been directed toward identifying MRI-derived characteristics in initially Aβ-negative individuals that may relate to future Aβ accumulation trajectories. Distinguishing such antecedent structural signatures from changes that emerge only after substantial Aβ deposition remains challenging, particularly because many MRI features associated with aging and AD overlap considerably. This makes it difficult to distinguish developmental predisposition from very early disease changes.

Our goal in this study was to understand whether cognitively healthy individuals exhibit early brain structural changes that predispose them to future Aβ accumulation and accelerated disease trajectory, using longitudinal data from the Fundació ACE Healthy Brain Initiative (FACEHBI). We focused on the analysis of cortical thickness, an indirect macrostructural proxy for synaptic loss that is considered to be more sensitive and specific to neurodegeneration than other measures like grey matter volume (Cai, et al., 2021; Kolasinski, et al., 2012; Ottoy, et al., 2025). Examining only those subjects that were not Aβ-positive at baseline, we found an association between brain structural anatomy and the future accumulation of Aβ pathology. This finding suggests that these anatomical patterns may reflect a latent vulnerability state preceding overt biomarker abnormality, rather than just downstream neurodegeneration.

## Results

### Differences in amount, velocity and acceleration of Aβ accumulation according to sex, MCI diagnosis and APOE genotype

Participants from the FACEHBI cohort underwent amyloid PET imaging at baseline (y0), two years (y2), and five years (y5). We examined Aβ burden at baseline and its rate of accumulation. Aβ burden was quantified through the CL in 125 Aβ-negative participants at baseline as a function of three groupings: sex, cognitive diagnosis — i.e. whether a mild cognitive impairment (MCI) diagnosis was endorsed at any follow-up assessment —, and genetic risk - i.e. whether the subjects had at least one APOE ε4 allele.

We detected that the CL score generally increased with age but was also subject to noise (**Figure 1A**). We observed CL group differences for cognitive diagnosis and APOE, but not for sex. Specifically, subjects who progressed to MCI during the study follow-up exhibited a higher baseline CL than subjects who remained as subjective cognitive decline (SCD) (**Figure 1B**, centre; age-corrected p-value=0.0008); although the rate of Aβ accumulation was not significantly different between them (**Figure 1C**, centre). With respect to APOE, subjects with at least one APOE ε4 allele exhibited both a higher baseline CL (**Figure 1B**, right; p-value<0.0001) and a faster accumulation than non-APOE ε4 carriers (**Figure 1C**, right; p-value=0.019). Finally, the accumulation of CL accelerated longitudinally for the entire cohort —i.e., the CL change from y2 to y5 was larger than the change from y0 to y2. This is represented in **Figure 1D** as a 2D joint distribution of the year change for the y0-y2 and the y2-y5 periods. No significant differences in CL acceleration between groups was found for any of the groupings (histograms not shown).

**Figure 1.**
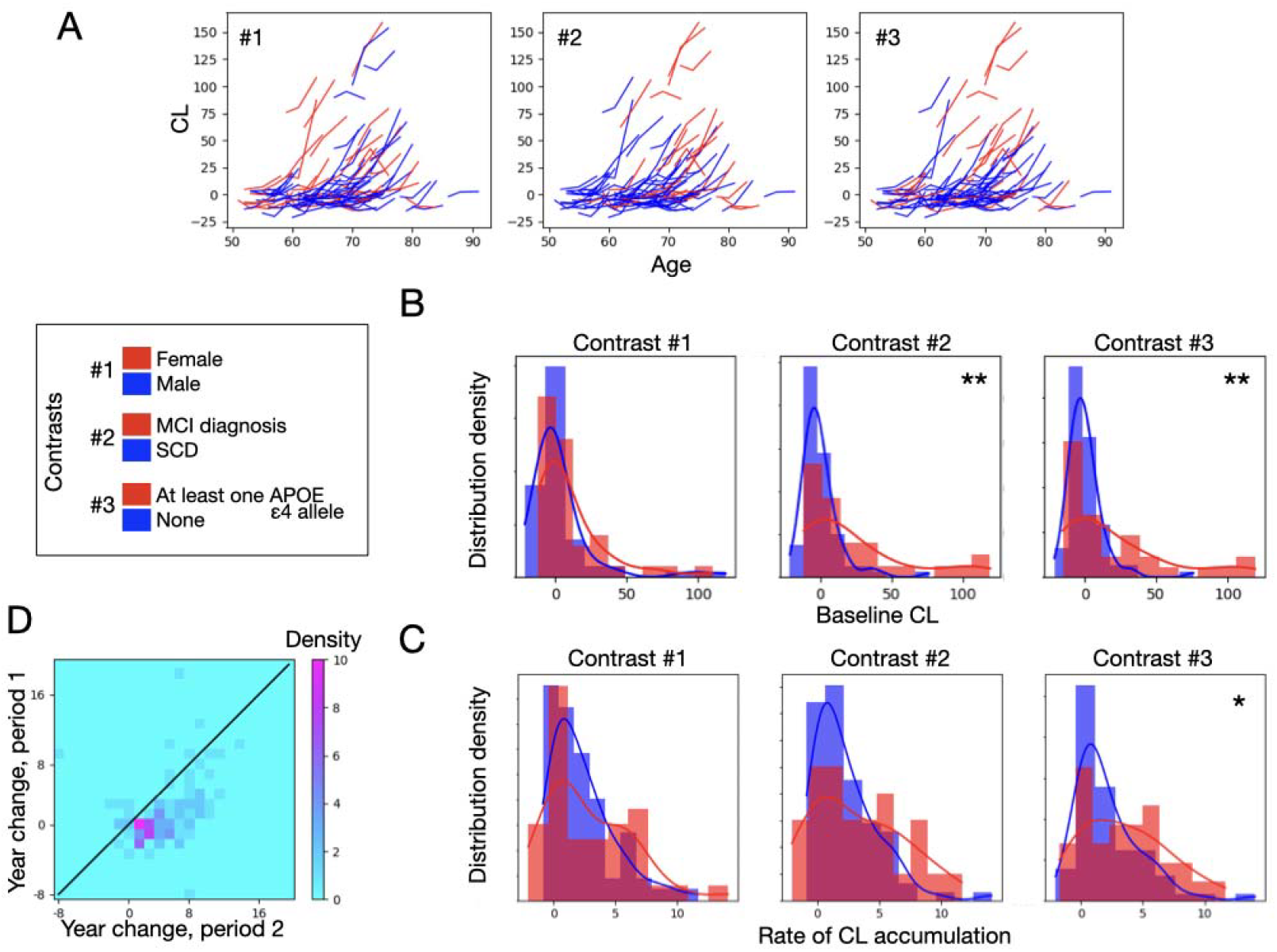
Aβ baseline and accumulation over the span of five years (visits in year 0, 2 and 5), characterised by sex (contrast #1), presence of an MCI diagnosis at any time during the five years (contrast #2), and genetic risk (having at least one APOE ε4 allele; contrast #3). **A**. Raw depiction of CL as a function of age, where each subject is represented by a curve. **B**. Distribution of the baseline CL per contrast; a positive diagnosis (MCI or dementia) and genetic risk (at least one APOE ε4 allele) correspond to statistically significant larger values. **C**. Distribution of rate of Aβ accumulation, with a significant effect of the APOE ε4 allele. **D**. Two-dimensional density of the year change for the first period (y0-y2) and the second period (y2-y5), showing an acceleration. (* Indicates p-value<0.05; ** indicates p-value<0.01).

For comparison, **Figure SI-1** shows the evolution in the cohort of two neuropsychological tests commonly used for diagnosis: MMSE and FNAME. These appear considerably noisier, and do not show a significant change on average from one session to the next.

### Cortical thickness at baseline vs. future rate of Aβ accumulation

Having characterised how Aβ accumulates over time in the FACEHBI cohort, we then focused on the main hypothesis of this study: whether the brain anatomy at baseline (y0), in terms of cortical thickness patterns across the cortex, showed an association with the rate of Aβ accumulation across 5 years. For these analyses we only included those subjects that were under a certain threshold of CL at y0. We varied this threshold from 0 to 20 to evaluate the sensitivity of the analysis to this parameter. The number of subjects selected per CL threshold level is shown in **Figure SI-2A** (from N=52 at the most restrictive threshold to N=101 at the other end). To obtain a degree of spatial granularity, we performed the analysis for each lobe separately: frontal, temporal, occipital, parietal, frontal and limbic (see **Figure SI-2B**, showing the number of ROIs per lobe). The following results were based on multivariate regressions that aimed to predict the CL increase from y0 to y5 from baseline brain MRI data (y0) —see **Methods**. Therefore, per CL threshold and lobe, we had one p-value (and a bootstrap-derived confidence interval).

We observed a strongly significant effect of cortical thickness to predict CL increase in the frontal lobe, and milder significant effects in parietal and limbic lobes that did not survive correction for multiple comparisons (correction across lobes). The effects in the frontal lobe decreased as we increased the CL threshold level, as subjects with possible incipient disease enter the sample. This is illustrated in **Figure 2**, which shows F-statistics (left) and (permutation-testing-based) p-values (right).

**Figure 2.**
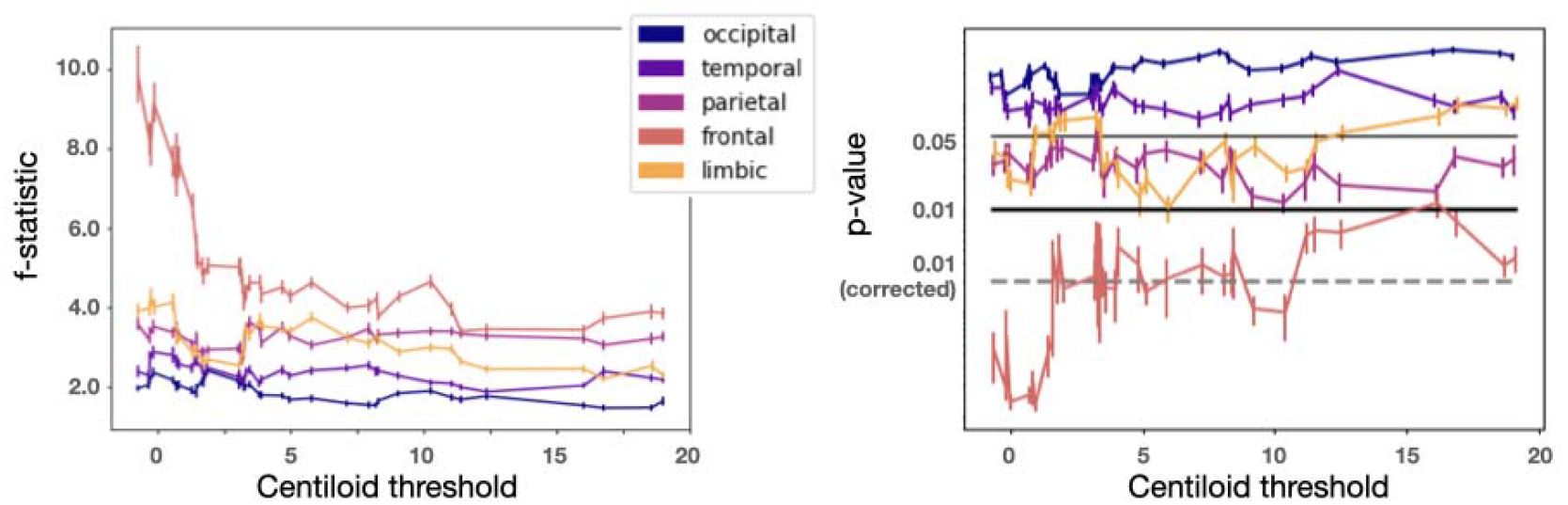
Relationship between cortical thickness at baseline and future rate of Aβ accumulation. This was instrumentalised as a multivariate regression, per lobe, of the rate of Aβ accumulation on the MRI values at y0. The regressions were run only on subjects that had no substantial Aβ at baseline —i.e. that were under a certain threshold of CL, which we varied across the range shown in the X-axis of the panels. **A**. F-statistics showing the strongest effect in the frontal lobe. **B**. Permutation-testing p-values; horizontal bars represent significance levels, with the dashed line representing Bonferroni-corrected significance level (correction across lobes). Error bars reflect statistical variability of the analyses in both panels (see **Methods**).

### Anatomical predisposition is spatially specific

Having identified a statistically significant correlation between baseline cortical thickness in the frontal lobe and subsequent rate of Aβ accumulation, we further examined the granularity of this pattern: whether this predisposition simply reflected overall reduced cortical thickness across the frontal lobe, or instead corresponded to a regionally specific pattern within frontal subregions. We addressed this question by sequentially perturbing the data, such that, in each perturbation, we broke a little of the spatial structure of the data within the frontal lobe (see **Methods** for details). As in the previous section, we then used multivariate regression to test the association between rate of Aβ accumulation and cortical thickness at baseline (here considered CL<20).

**Figure 3A** shows the regression coefficients (or betas) for each frontal region of interest for the original (before perturbation) data. This shows a distributed mixture of positive and negative values, suggesting, in principle, a fine-grained pattern across regions (see **Table SI-1**). **Figure 3B** presents one curve of explained variance for repetition of the perturbation procedure, showing a decline in model performance as we perturbed the matrix. This result argues for a spatially-specific pattern (where regions are not exchangeable) and against a broad, nonspecific association with full-lobe cortical thickness. **Figure 3C** shows the percentage of statistically significant predictions (from 100 bootstrapped repetitions of the analysis) as we keep adding perturbations on the data. The sharp decrease offers quantitative evidence in favour of the result.

**Figure 3.**
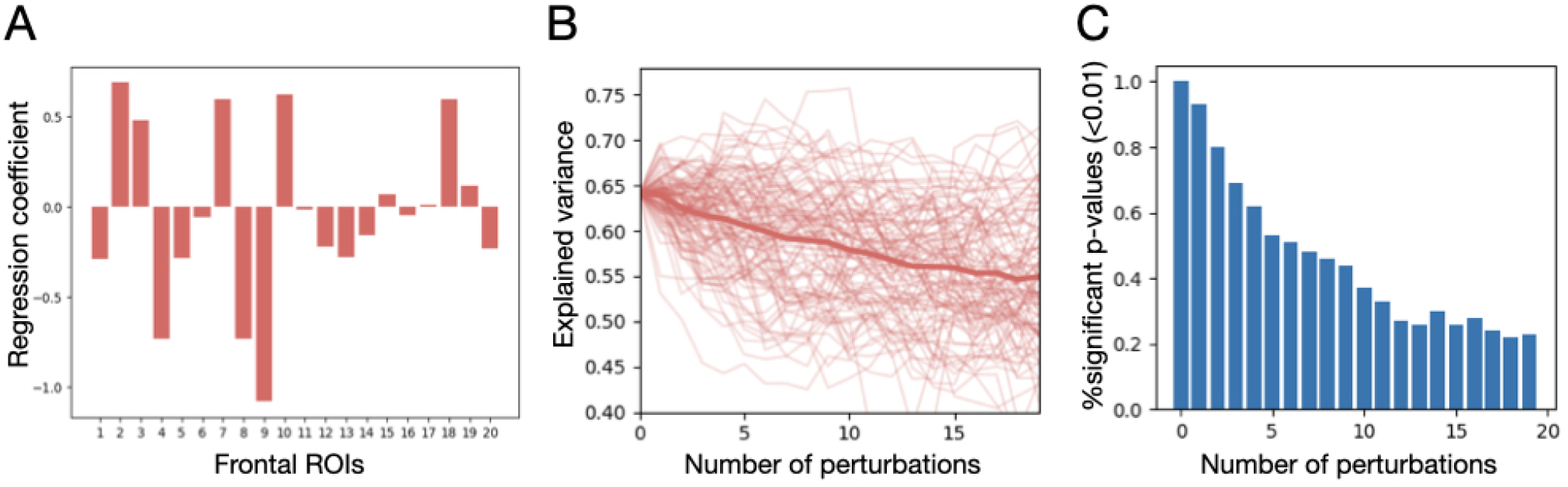
Spatial-granularity of the frontal lobe predisposition pattern. **A**. Regression coefficients (betas) from regression future rate of Aβ accumulation on frontal cortical thickness at baseline; suggesting a spatial-specific pattern, the betas are positive and negative, and widely distributed (**Table SI-1** contains the list of frontal ROIs). **B**. To quantitatively assess the spatial specificity of the prediction, we sequentially induced perturbations on the data by swapping the values of randomly chosen pairs of regions for a random subset of the subjects. We repeated the procedure 100 times. The curves reflect the loss in explained variance as we perturb the data; the average is represented by the thicker line. **C**. Percentage of statistically significant predictions (out of 100) for each model as we perturb the data.

### Cortical thickness at baseline vs. APOE

Previous results showed the association between APOE and Aβ accumulation in the study cohort, as well as between baseline cortical thickness and Aβ accumulation. We next examined the relation between APOE and cortical thickness at baseline. Here, our regression model aimed at predicting baseline cortical thickness for each region of interest from a design matrix with five dummy variables representing the available genetic profiles: ε2/ε3, ε3/ε3, ε2/ε4, ε3/ε4 and ε4/ε4 (which N in the full cohort is, respectively: 18, 71, 3, 32 and 2). As before, the regression was performed for each threshold level of CL —see **Methods** for details.

Figure 4. shows the p-values per region, coloured per lobe (one dot per region); together with the geometric mean across regions (represented by lines). Primarily, only some regions in the parietal cortex turned significant, but none survived correction for multiple comparisons via Bonferroni (correction across regions; threshold out of range and not shown).

## Discussion

In the classical model of AD progression, Aβ deposition represents the earliest detectable event and is believed to trigger subsequent pathological changes, including production of hyperphosphorylated tau aggregates and neuronal death, which together lead to cognitive decline (Hardy & Higgins, 1992). Amyloid accumulation thus precedes detectable brain atrophy (Xia, et al., 2025). While the causal role of Aβ in AD remains controversial (Granzotto & Sensi, 2024), it is a widely used biomarker (Do-Hoon, 2025) and is associated with future cognitive decline (Gallego-Rudolf, Wiesman, Binette, Villeneuve, & Baillet, 2024). In this study, we investigated the brain anatomical patterns of cortical thickness that predispose to Aβ accumulation before the onset of deposition. For this purpose, we used data from a longitudinal cohort of individuals with subjective cognitive decline, focusing on those with no evidence of Aβ deposition at baseline. The temporal dynamics of Aβ deposition and their statistical relationship with APOE and diagnosis in the cohort were in line with the literature (Bollack, et al., 2024).

**Figure 4.**
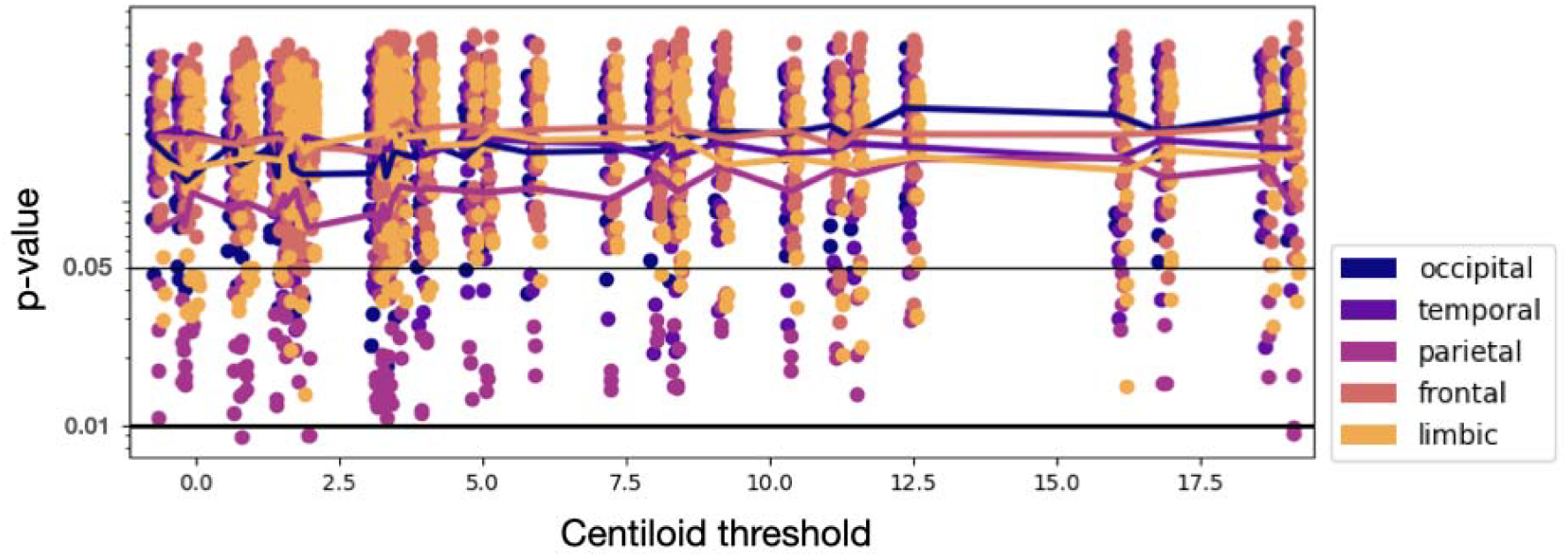
Relation between APOE and cortical thickness at baseline, quantified as a multivariate regression of each region’s cortical thickness at baseline on a design matrix reflecting all available APOE profiles. The analysis was run only on those subjects that were under a certain CL threshold at baseline; we varied the threshold for a range of values up to 20. Each point represents a region’s p-value, coloured by lobe. Horizontal lines reflect significance levels; Bonferroni-corrected (across-regions) significance level is below the range and not shown.

Our main result demonstrates a significant association between baseline frontal cortical thickness and the subsequent rate of Aβ accumulation, suggesting the presence of an early anatomical predisposition fingerprint within the frontal lobe for Aβ deposition and later development of AD, prior to the emergence of other signs. No clear association between these baseline cortical thickness patterns and APOE genotype could be identified, even though APOE was previously shown in this same cohort to be associated with both early Aβ accumulation and cognitive performance (Moreno–Grau, et al., 2018). This suggests that APOE and cortical thickness fingerprints may be two dissociable causes of amyloid accumulation. Importantly, this predisposition could not be described as a broad decrement of cortical thickness (i.e. it could not be summarized with a single value), but as fine-grained cortical patterns reflecting the relative value of cortical thickness across areas. While early anatomical changes have been reported prior to the onset of AD symptoms (Smith, Chebrolu, Wekstein, Jicha, & & Markesbery, 2007; Tondelli, et al., 2012), these changes are primarily observed in temporal and hippocampal areas, although certain specific frontal regions (such as the orbitofrontal cortex) have been also implicated (Tondelli, et al., 2012). Although the frontal cortex is well-recognised to play a central role in memory encoding and consolidation (Frankland, 2005; Kitamura, et al., 2017), further work is required to determine whether the frontal fingerprint of AD susceptibility that we observe here has any associated behavioural correlate and how, if at all, it is causally related to the subsequent development of pathology.

Related to this work, distinct patterns of anatomical change in preclinical subjects were found depending on the presence of Aβ at baseline (Pegueroles, et al., 2017). Our question is however complementary: how baseline anatomic structure influences subsequent Aβ accumulation? This type of question and analysis help to separate anatomical predisposition from early disease changes. Notably, the association between brain structural changes and subsequent Aβ accumulation is only detectable when the analyses are restricted to subjects with low or absent baseline Aβ burden (using a set of CL thresholds up to 20). One possible explanation is that, indeed, as the CL threshold is increased and additional participants are included, the analysis increasingly reflects a mixture of two different processes: predisposition effects and early AD stages. These processes may be associated with different anatomical signatures, rendering a simple linear regression model, particularly in a small sample, inappropriate.

Our results align with data from animal models that show a connection between baseline brain structure and future amyloid accumulation. For instance, in a transgenic mouse model of AD, structural differences were identified prior to the onset of amyloid formation and predicted where amyloid would accumulate at a later point in time (Badea, Johnson, & Jankowsky, 2010). Importantly, plain amyloid mouse models (i.e. without tau) do not often show atrophy, suggesting that these morphological differences are unlikely to reflect a neurodegenerative process that already started, but a pre-existing structural trait (Kuhla, et al., 2017).

In conclusion, these results indicate that certain patterns of brain morphology may confer increased susceptibility to the onset of early changes in the AD cascade of events. These morphological signatures may reflect genetic predisposition, lifestyle factors or a combination of both. Despite the moderate sample size, which should be expanded to attain clinically useful predictive power, our findings are important because they may enable better identification of individuals at risk of AD and contribute to the understanding of the biological sequence of disease progression.

## Methods

### Study cohort

FACEHBI is a single-centre longitudinal study of aging, cognition, lifestyle and biomarkers from 200 individuals with Subjective Cognitive Decline (SCD) conducted at Ace Alzheimer Center Barcelona from 2014 (Rodriguez-Gomez, et al., 2017). The inclusion criteria are detailed elsewhere, but, in short, they comprised: age ≥ 50 years; presence of subjective cognitive complaints (defined as a score of ≥ 8 on MFE-30, the Spanish version of the Memory Failures in Everyday Life Questionnaire (Lozoya-Delgado, Ruiz-Sánchez de León, & Pedrero-Perez, 2012)) but objectively normal cognition on a comprehensive neuropsychological battery; a Mini-Mental State Examination (MMSE) score ≥ 27, and no functional impairment (Clinical Dementia Rating = 0). Additional criteria included at least six years of formal education, no major depression or anxiety (HADS < 11), no history of epilepsy or significant renal or hepatic disease, and no substantial sensory impairments. Participants were evaluated yearly with neurological and cognitive exams. Additionally, at baseline visit (y0) and follow-up visits 2 (y2) and 5 (y5), several biomarkers, including brain MRI and an amyloid-PET, were performed in a 90-day window after clinical evaluations.

Written consent was obtained from all participants prior to the enrolment in the study. The FACEHBI protocol received approval from the ethics committee of Hospital Clínic i Provincial in Barcelona, Spain (EudraCT number 2014–00079-38). Collection protocols were in accordance with ethical standards according to World Medical Association Declaration of Helsinki—Ethical Principles for Medical Research Involving Human Subjects.

In the present study, data from 125 participants who underwent three longitudinal amyloid-PETs and brain MRI scans were analysed.

### Cognitive evaluation

Participants underwent a neuropsychological protocol that included the Mini-Mental State Examination (MMSE) (Folstein, Folstein, & McHugh, 1975; Blesa, et al., 2001), the neuropsychological battery of Fundació ACE (NBACE) (Alegret M., et al., 2012; Alegret M., et al., 2013), the Spanish version of the Face Nname Associative Memory Exam (S-FNAME) (Amariglio, et al., 2012; Alegret, et al., 2015), and a set of self-administered questionnaires (Rodriguez-Gomez, et al., 2017). All participants had a diagnosis of SCD at baseline and their neurological and cognitive status was reassessed yearly, assessing their potential conversion to mild cognitive impairment (MCI). A diagnosis of MCI was endorsed if performance in any of the NBACE test scores was impaired according to the published cutoffs and participants maintained their autonomy in daily life activities (Alegret M., et al., 2012; Alegret M., et al., 2013).

### Brain MRI

Structural MRI data were acquired for all participants prior to PET scanning. MRI was performed at the Clínica Corachan, Barcelona on a Siemens Magnetom Aera 1.5 T scanner (32-channel head coil) using a T1-weighted 3D MPRAGE sequence (TR = 2200 ms, TE = 2.66 ms, TI = 900 ms, flip angle = 8°, field of view = 250 mm, slice thickness = 1 mm, isotropic voxels 1 × 1 × 1 mm). Additional sequences included axial T2-weighted, 3D FLAIR, and T2*-weighted acquisitions for detection of vascular pathology or microbleeds. Preprocessing steps included: visual inspection for artefacts and manual centring to the anterior commissure; segmentation of T1 images into grey matter (GM), white matter (WM), and cerebrospinal fluid (CSF) using SPM12 and DARTEL to normalise to MNI space; calculation of total intracranial volume (TIV) as GM + WM + CSF; smoothing of GM images with an isotropic Gaussian kernel (e.g., 6 mm FWHM) for voxel-based morphometry (VBM); extraction regional cortical thickness using FreeSurfer (Cardinale, 2014).

After a visual inspection for artifacts, cortical and subcortical segmentation of the structural images was performed using FreeSurfer 7.4 (https://surfer.nmr.mgh.harvard.edu/). This procedure allows the segmentation of gray matter, white matter, and other substructures (Fischl B. FreeSurfer. Neuroimage. 2012;62:774–81.).

### Amyloid-PET

Aβ PET imaging used the tracer 18F-Florbetaben (NeuraCeq). Scans were acquired on a Siemens Biograph PET/CT system at the Radiology Department from the Hospital Clínic i Provincial in Barcelona. Four 5-minute frames were collected beginning approximately 90 minutes after intravenous injection of about 300 MBq of tracer. PET images were co-registered to the participant’s structural T1 MRI, motion-corrected, averaged, and normalised to standard space. Standard uptake value ratios (SUVR) were computed as the mean uptake in a composite cortical region (frontal, temporal, parietal, and posterior cingulate cortices) divided by uptake in the cerebellar cortex as reference. SUVRs were converted to Centiloid (CL) units following the standard Centiloid transformation procedure (Klunk, et al., 2015; Kolinger, et al., 2025).

### APOE genotyping

Genomic DNA was extracted from peripheral blood using the commercially available Chemagic system (Perkin Elmer). APOE genotypes were extracted from the Axiom SP array (Thermo Fisher Scientific) (Moreno-Grau, et al., 2019). Ace Alzheimer Center Barcelona has this variable as a standard in its assessment protocols. Alternatively, the APOE genotypes were determined using fluorogenic allele-specific oligonucleotide probes (TaqMan assay; Life Technologies, Spain) for rs7412 (Test ID: C 904973_10) and rs429358 (Test ID: C 3084793_20). For the TaqMan assays, PCR and real-time fluorescence measurements were carried out on a QuantStudio3 real-time PCR system (Thermo Fisher Scientific, Spain) using the TaqMan Universal Master Mix (ref: 4364341, Life Technologies, Spain) methodology according to manufacturer’s instructions. The polymerase chain reaction was performed as follows: first, a pre-read step for 30 s at 60C, denaturation for 10 min at 95C, followed by 40 cycles at 95C for 15 s and 60 C for 1 min, and a post read stage for 30 s at 60C. The genotype was determined using the Genotyping App for Thermo Fisher Scientific Cloud by clustering analysis. The laboratory technicians were blinded to other study variables.

### Statistical analyses

We performed four sets of analyses. First, using the CL as a metric of Aβ deposition, we quantified differences in the average Aβ deposition and its rate accumulation between groups given by (i) sex, (ii) cognitive diagnosis (conversion to MCI during the follow-up), and (iii) APOE genotype (presence of at least one APOE ε4 allele). The rate of accumulation was defined as the slope from (linearly) regressing CL on the subject’s age at the time of the PET scan. Statistical significance of the differences between groups was assessed using permutation testing. This analysis corresponds to **Figure 1**.

Second, we assessed the relation between the rate of Aβ accumulation (quantified as before), and MRI-derived cortical thickness at baseline (y0). Having calculated cortical thickness for each ROI, we performed the analyses per cerebral lobe (frontal, temporal, occipital, parietal, frontal and limbic) to have some extent of spatial granularity — see **Figure SI-2B**, showing the number of ROIs per lobe. Thus, for each lobe, we ran a multivariate regression model where we predicted the estimated the rate of Aβ accumulation from the MRI-derived cortical thickness values at baseline (y0). We controlled by age at baseline, gender and intracranial volume by regressing these confounds out from both the dependent and independent variables. This multivariate regression model was embedded into a permutation testing procedure (10001 permutations) to compute a p-value per model. Importantly, to be able to select subjects that are truly healthy at baseline, we used only these subjects that are under a certain threshold of CL at y0. Typically, CL values equal or lower than 10 or less are considered indicative of absence of amyloid pathology (Amadoru, et al., 2020), whereas values larger than 26 are indicative of significant amyloid deposition, consistent with AD pathology, and a clear predictor of progression to dementia (Hanseeuw, et al., 2021). Here, we varied this threshold from ~0 to ~20 to evaluate the sensitivity of the analysis to this parameter (see **Figure SI-2A**, showing the number of subjects per threshold level). To quantify the uncertainty of our estimations, for each lobe and CL threshold level, we embed the entire analysis within a bootstrap scheme (100 bootstrap repetitions with resample of subjects with repetition). Altogether, this procedure returned a (lobes–by–thresholds) matrix of effect sizes (f-statistics) and p-values per bootstrap repetition. We took the average to estimate expected means and computed non-parametric confidence intervals (1%-99%). This is the analysis used for **Figure 2** (where we show f-statistics instead of standard effect sizes like explained variance, because in-sample explained variance depends on the number of features and data points, which varies across lobes and thresholds).

Having found a significant effect on the frontal lobe, the third analysis aimed at quantifying the granularity of this pattern in order to discern whether this was a broad effect owing to generally reduced cortical thickness across the entire frontal lobe, or a specific pattern across frontal regions. To answer this question, we sequentially perturbed the data such that, for each perturbation, we randomly chose two regions and swapped the cortical thickness values between the two regions for (a randomly chosen) half of the cohort. The cortical thickness matrix was previously standardised across subjects such that there was no difference between regions across the population. We then ran multivariate regressions of rate of Aβ accumulation on the cortical thickness values after each perturbation and assessed statistical significance using permutation testing. We repeated the process 100 times. This is the analysis used for **Figure 3**.

Finally, we assessed the relation between genetics and frontal cortical thickness at baseline. For this, we created a design matrix with as many rows as subjects and five columns that correspond to the available genetic profiles: ε2/ε3, ε3/ε3, ε2/ε4, ε3/ε4 and ε4/ε4. We then used this design matrix to predict cortical thickness at baseline for each region of interest. As before, the regression was performed for each threshold level of the CL and embedded in a permutation testing scheme. The entire procedure was bootstrapped 100 times; to combine across the 100 p-values, we used the Fisher combination. This is the geometric mean, i.e. exponentiating the average of the logarithm of the 100 p-values —a combination technique that amplifies the importance of values near zero (Fisher, 1932; Vidaurre, et al., 2019). This analysis was used in **Figure 4**.

## Acknowledgements

We are grateful to all FACEHBI participants, without whom this study would not have been possible. We thank all FACEHBI sponsors for making this possible and all of the investigators from Ace Alzheimer Center Barcelona, Hospital Clinic i Provincial de Barcelona, Clínica Corachan and Life Molecular Imaging GmbH for their close collaboration and continuous intellectual input. The FACEHBI study group includes:

Aguilera N^1^, Alarcón-Martín E^1^, Alegret M^1,2^, Alllué JA^3^, Bayón-Bujan P^1^, Berthier M^4^, Boada M^1,2^, Buendia M^1^, Bullich S^5^, Campos F^6^, Calm-Salvans B^1^, Cano A^1,2^, Casales F^1^, Cañabate P^1,2^, Cañada L^1^, Cuevas C^1^, de Rojas I^1^, Diego S^1^, Domingues-Kolinger G^5^, Escudero JM^7^, Espinosa A^1,2^, Fandós N^3^, Fernández MV^1^, Gailhajenet A^1^, García-González P^1,2^, Giménez J^7^, Gomez-Chiari M^7^, Guitart M^1^, Gurruchaga MJ,^1^ Gutiérrez PC,^1^ Hernández I^1,2^, Ibarria M^1^, Lafuente A^1^, Lomeña F^6^, Marquié M^1,2^, Martín E^1^, Martínez C,^1^ Martinez M,^1^ Miguel A,^1^ Moreno M^1^, Morera A^1^, Montrreal L^1^, Muñoz N^1^, Muñoz-Morales A^1^, Niñerola A^6^, Nogales AB^1^, Núñez L^8^, Olivé C^1^, Orellana A^1,2^, Ortega G^1,2^, Páez A^8^, Pancho A^1^, Pelejà E^1^, Pérez-Martínez E^5^, Pérez-Cordon A^1^, Pérez-Grijalba V^3^, Pascual-Lucas M^3^, Perissinotti A^6^, Preckler S^1^, Puerta R^1^, Ramis MI^1^, Roé-Vellvé N^5^, Romero J^3^, Rosende-Roca M^1^, Ruiz A^1,2^, Sanabria A^1,2^, Sanz-Cartagena P^1^, Sarasa L^3^, Seguer S^1^, Seimandi C,^1^ Solivar A,^1^ Sotolongo-Grau O^1^, Stephens A^5^, Tartari JP^1^, Tárraga L^1,2^, Tejero MA^7^, Terencio J^3^, Torres M^8^, Valenzuela A,^1^ Valero S^1,2^, Vargas L^1^, Vivas A^7^.

^1^ Ace Alzheimer Center Barcelona – Universitat Internacional de Catalunya (UIC). Barcelona, Spain

^2^ CIBERNED, Center for Networked Biomedical Research on Neurodegenerative Diseases, National Institute of Health Carlos III, Ministry of Economy and Competitiveness. Madrid, Spain

^3^ Araclon Biotech-Grifols. Zaragoza, Spain

^4^ Cognitive Neurology and Aphasia Unit (UNCA). University of Malaga. Málaga, Spain

^5^ Life Molecular Imaging GmbH. Berlin, Germany

^6^ Servei de Medicina Nuclear, Hospital Clínic i Provincial. Barcelona, Spain

^7^ Departament de Diagnòstic per la Imatge. Clínica Corachan, Barcelona, Spain

^8^ Grifols®. Barcelona, Spain

Part of the data used in the preparation of this article were obtained from the Prognostic and Natural History Study (PNHS), provided by the Amyloid Imaging to Prevent Alzheimer’s Disease Consortium (AMYPAD). As such, investigators within the AMYPAD PNHS and AMYPAD Consortium contributed to the design and implementation of AMYPAD and/or provided data but did not participate in the analysis or writing of this report.

AMYPAD consortia representative: Prof. Frederik Barkhof, Department of Radiology and Nuclear Medicine, Amsterdam UMC, Vrije Universiteit Amsterdam, The Netherlands.

A complete list of AMYPAD investigators can be found at https://doi.org/10.5281/zenodo.7962737. A complete listing of AMYPAD contributors can be found at: https://amypad.eu/partners/.

## Supplementary Information

**Figure SI-1.**
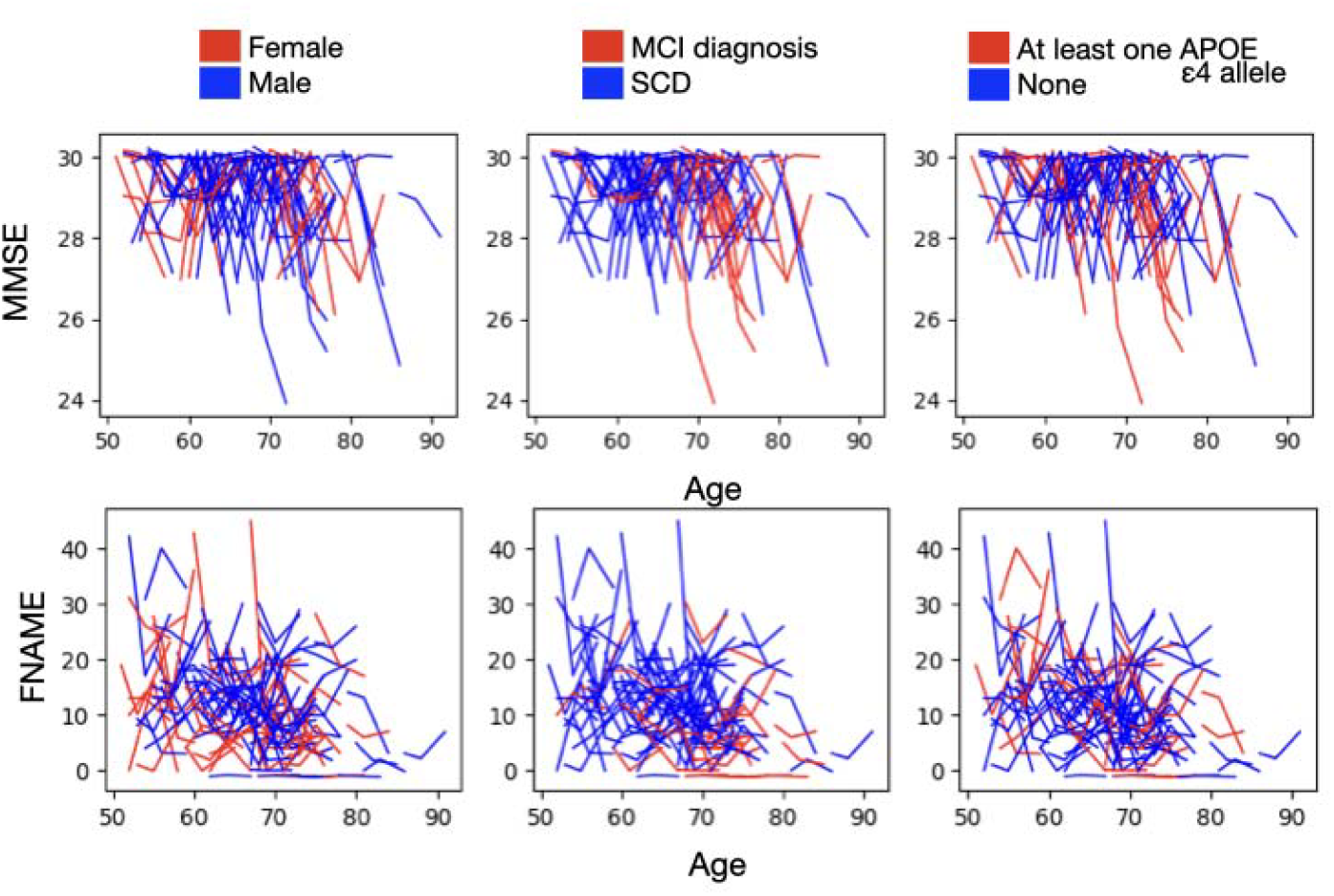
Cognitive scores, as measured by MMSE and FNAME, as a function of age, for three different contrasts: sex, diagnosis and APOE; each subject is represented by a curve.

**Figure SI-2.**
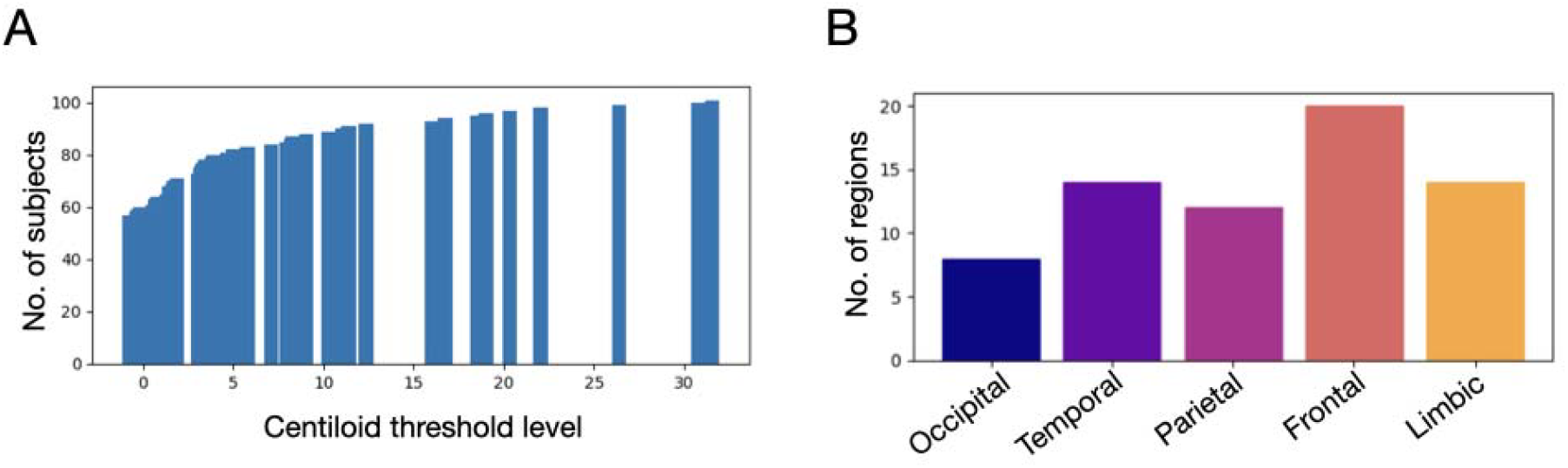
**A**. Number of subjects after discarding those that were above a CL threshold level, across different thresholds. **B**. Number of regions of interest per lobe.

**Table SI-1.** List of frontal ROIs.

| ROI number | Description |
| --- | --- |
| 1 | caudal middle frontal left |
| 2 | frontal pole left |
| 3 | lateral orbitofrontal left |
| 4 | medial orbitofrontal left |
| 5 | pars opercularis left |
| 6 | pars orbitalis left |
| 7 | pars triangularis left |
| 8 | precentral left |
| 9 | rostral middle frontal left |
| 10 | superior frontal_left |
| 11 | caudal middle frontal right |
| 12 | frontal pole right |
| 13 | lateral orbitofrontal right |
| 14 | medial orbitofrontal right |
| 15 | pars opercularis right |
| 16 | pars orbitalis right |
| 17 | pars triangularis right |
| 18 | precentral right |
| 19 | rostral middle frontal right |
| 20 | superior frontal right |

